# iMyocyte: A Web-Based Cellular Automata Platform for Real-Time Demonstration of Cardiac Reentry

**DOI:** 10.64898/2026.02.02.703157

**Authors:** Lareina Shen, Phoenix Plessas-Azurduy, Gil Bub

## Abstract

Cardiac arrhythmias arise from excitable-media dynamics that are often challenging to teach and visualize in real time. To address this, we developed *iMyocyte*, an interactive web-based platform for real-time demonstration of excitation wave propagation, conduction block, and reentry dynamics using a simplified cellular automaton model (i.e., a network of discrete cells that change state based on neighbouring cells). In *iMyocyte*, a network is built from student devices, each representing a model cell coupled to its neighbours to propagate excitation. Unlike traditional teaching materials, *iMyocyte* lets students manipulate key model parameters and immediately observe how these shape wave dynamics and reentry conditions. Pilot classroom deployments with undergraduate life science students at McGill University indicate high engagement and suggest improved conceptual clarity, motivating future work to improve the platform’s usability, robustness, and visualization features, and to conduct formal learning assessment.

## 1 Introduction

Cardiac arrhythmias and sudden cardiac death represent a major global health problem, accounting for approximately 15–20% of all deaths [1]. Many clinically important arrhythmias, including atrial fibrillation, atrioventricular (AV) tachycardia, and ventricular tachycardia, are sustained by reentry [2]. Reentry occurs when an activation wavefront circulates around a pathway and repeatedly re-excites recovered excitable tissue, producing rapid, self-sustaining rhythms (Figure 1) [3, 4]. The reentry concept traces to George Mines’ early 20th-century work demonstrating that excitation can perpetuate in closed loops when conduction and recovery timing permit a returning wavefront to encounter excitable tissue [3].

**Figure 1:**
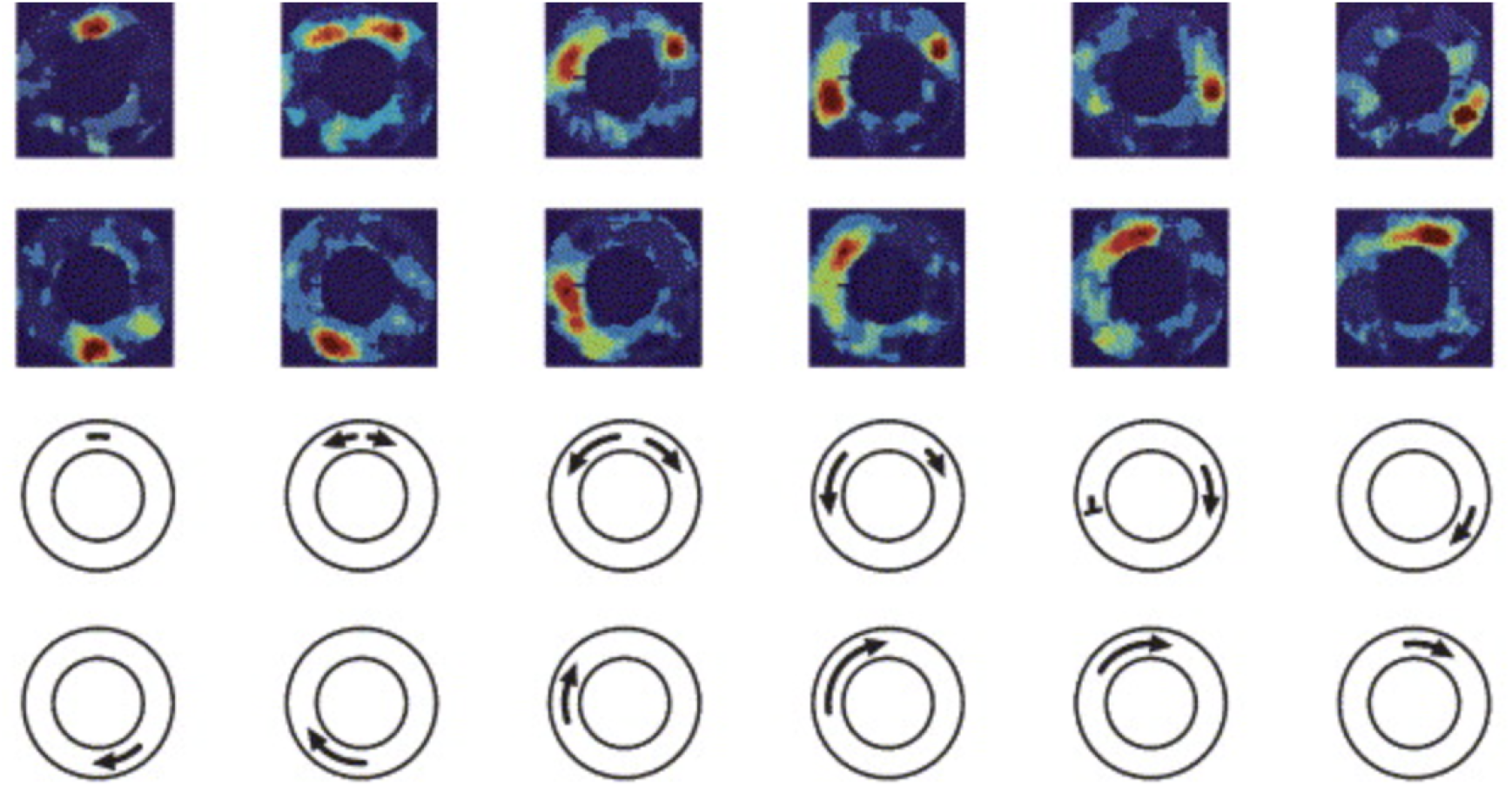
Illustrative visualization of reentry in excitable media. Unidirectional block can initiate a circulating wave that sustains reentry when it returns after local recovery (adapted from [10]).

Although reentry is well characterized mechanistically, building intuition for its spatiotemporal evolution can be challenging when instruction relies primarily on static diagrams. Mechanistic computational models provide a reproducible framework for probing how specific parameter changes alter cardiac dynamics and have become central tools in cardiac electrophysiology research [5]. However, high-fidelity, multi-scale simulations often require specialized software and substantial computational resources, limiting their suitability for routine, interactive use outside specialized research or clinical settings [6]. In contrast, simplified excitable-media models can reproduce key qualitative phenomena (e.g., wave propagation, conduction block, and reentrant spiral-wave dynamics) at much lower computational cost [6, 7]. These properties make such models a practical substrate for interactive tools aimed at developing intuition for reentry dynamics.

Interactive computational tools can make nonlinear excitable-media dynamics tangible by allowing learners to manipulate parameters and immediately observe emergent behaviour. Education research suggests that active, interactive learning can improve conceptual understanding and retention relative to purely didactic approaches [8]. Here, our goal is not to replace biophysically detailed simulators, but to provide an accessible, real-time demonstrator of core reentry principles suitable for classroom use, with explicit assumptions and limitations.

This study introduces *iMyocyte*, a multi-user web platform for real-time demonstration of wave propagation, conduction block, and reentry. We modeled *iMyocyte* after a cellular automaton (CA): a network of discrete cells whose state updates depend only on neighbouring cells, enabling low-cost simulation of qualitative wave dynamics [9]. By combining browser-based computation with publication/subscriber (pub/sub) coordination, *iMyocyte* lowers the barrier to hands-on, classroom-scale demonstrations without specialized hardware. Users can adjust key model parameters to explore how these shape wave dynamics and the conditions for reentry. We describe the model and distributed architecture, report pilot classroom deployments, and discuss scalability constraints and design improvements for robust use in larger cohorts.

### 2 Methods

### 2.1 Systems Overview

*iMyocyte* is a web-based platform in which each participant’s device runs a local instance of a simplified excitable-media CA. The system is designed so that (i) computation of cell-state updates occurs locally on each client, and (ii) the network layer is used to coordinate topology formation and to broadcast discrete state-change events (e.g., stimulation pulses and parameter updates) needed for consistent visualization across devices. Figure 2 summarizes the decentralized architecture of *iMyocyte*, in which client-side CA execution is coordinated via pub/sub messaging with run logging outside the simulation loop. While the platform supports multiple interaction modes, this study evaluates only the ring-topology workflow (implemented as “Neighbour Ring Mode”), which enables reproducible reentry demonstrations. The full source code and a runnable version of *iMyocyte* is publicly available on GitHub at github.com/lareinashen/iMyocyte, allowing reviewers and instructors to download, deploy, and interact with the platform directly.

**Figure 2:**
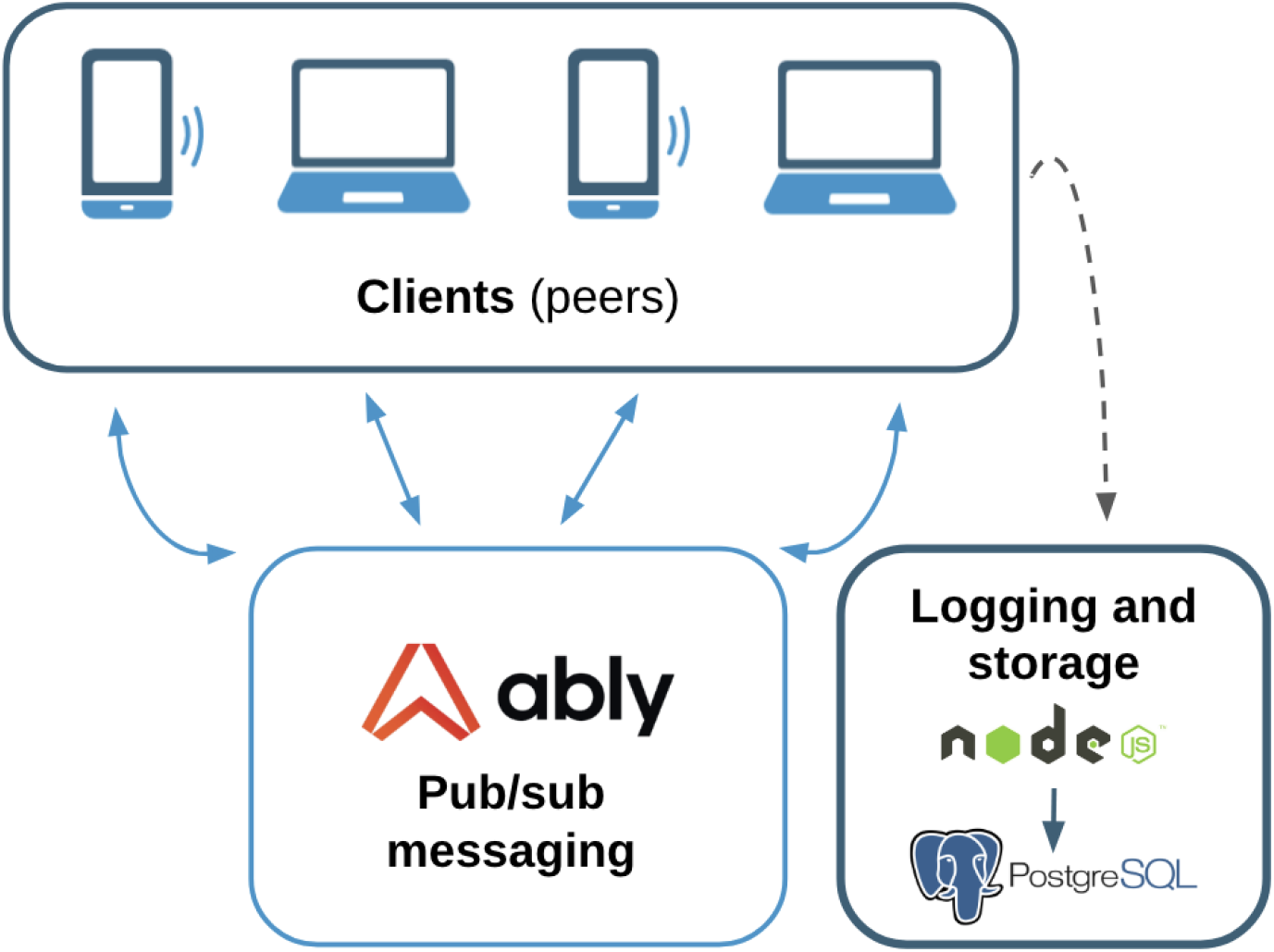
High-level *iMyocyte* architecture. Each participant device executes CA updates and renders visualization locally. Real-time coordination across devices (presence, topology/setup, stimulation events, and parameter updates) is mediated via WebSocket pub/sub messaging (Ably). A separate backend logging pipeline (Node.js →PostgreSQL) records run metadata for post hoc analysis and is not required for simulation execution.

#### 2.1.1 Client Implementation

The client application is implemented in React (v18.3.1) and TypeScript (v5.6.2), built with Vite (v6.0.5) [11–13]. React manages the user interface and visualization updates, while TypeScript enforces static typing to reduce runtime errors in a multi-user, real-time setting. Each client maintains (i) its local CA state (e.g., cell activity and health status), (ii) a record of its current neighbors, and (iii) CA timing parameters derived from a local JSON configuration file containing base values of conduction velocity (CV) and refractory period (RP) and exercise-dependent rate multipliers that vary by ring size, sympathetic activation mode, exercise level, and health status. All values are treated as relative scaling parameters rather than biophysically calibrated measures.

#### 2.1.2 Real-time Messaging and Session Coordination

Real-time communication between clients is handled by a commercial pub/sub service (Ably Realtime LTD; v2.6.2) [14]. The free tier is sufficient for the examples in this publication. Ably provides presence detection to identify active participants, message ordering within channels, and automatic reconnection after transient network interruptions. The platform leverages these features to (i) form and modify user-defined interaction graphs (e.g., ring topologies for reentry demonstrations) and (ii) propagate discrete state-change events across the network. Because network conditions can vary, the application treats the messaging layer as best-effort coordination rather than a guarantee of strict real-time determinism. Accordingly, the model is intentionally designed to be robust to small timing jitter during classroom use.

#### 2.1.3 Run Logging and Data Storage

During runtime, a backend API (Node.js) records run metadata for post hoc analysis (e.g., pseudonymous participant identifiers, topology snapshots, and timestamps of firing and reentry events). In our implementation, these records are stored in a PostgreSQL database. This logging pipeline is not part of the simulation loop: CA rule execution and real-time coordination occur client-side, and the simulation can continue to run even if the logging service is unavailable.

### 2.2 Cellular Automaton Model

*iMyocyte* models excitable tissue as a network of discrete cells coupled by local interactions. Each cell, *i*, follows a three-state excitable-media CA:

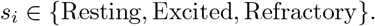

This model is inspired by the Greenberg–Hastings framework, a minimal excitable-media CA that reproduces qualitative wave propagation and conduction block using discrete state transitions [15]. An incoming activation excites a cell only if it is Resting; activations arriving during Excited or Refractory are ignored (discrete conduction block). When a cell becomes Excited, it (i) schedules activation events to its neighbours after a propagation delay and (ii) schedules its own recovery to Resting after a RP.

#### Propagation delay

For each cell-cell connection (*i, j*) in the ring, activation is delivered after a fixed propagation delay *τ*_*i*_, where *τ*_*i*_ denotes the event latency from cell *i* to its neighbour. The delay is computed as

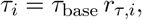

where *τ*_base_ sets the apparent wave speed and *r*_*τ,i*_ is a cell-specific multiplier. In the implementation, the user-facing “CV” control is an inverse proxy for speed: decreasing *τ*_*i*_ produces faster apparent conduction, so we report propagation delay *τ*_*i*_ (ms) rather than physical conduction velocity (m/s).

#### Refractory period

After a cell fires, it enters a refractory state for duration 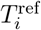, during which it cannot be re-excited. This duration is computed as

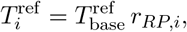

where 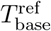 is a baseline refractory period and *r*_*RP,i*_ is a cell-specific multiplier.

#### Parameter modulation

Multipliers *r*_*τ,i*_ and *r*_*RP,i*_ are determined by a cell’s health status, the exercise level, and presence of global sympathetic activation. Dynamic changes to these multipliers alter the balance between wave propagation speed and cell recovery, thereby affecting reentry propensity.

As shown in Table 1, higher exercise intensity and global sympathetic activation decrease propagation delay and shorten RP in the *iMyocyte* model. This reflects the qualitative effects of increased sympathetic ( -adrenergic) drive during exercise, which typically shortens effective refractory periods and accelerates cardiac activation dynamics in support of increased heart rate and contractility during exertion [16–18]. Unhealthy cells in *iMyocyte* are assigned slower propagation and longer RP to represent pathological substrates (e.g., fibrosis or ischemic injury) that slow conduction and increase recovery heterogeneity [16, 19].

**Table 1:**
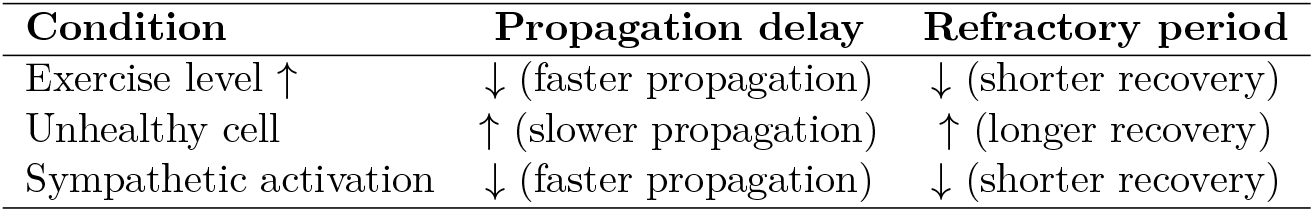
Qualitative parameter modulation used in *iMyocyte*. Arrows indicate the direction of change relative to baseline.

| Condition | Propagation delay | Refractory period |
| --- | --- | --- |
| Exercise level ↑ | ↓ (faster propagation) | ↓ (shorter recovery) |
| Unhealthy cell | ↑ (slower propagation) | ↑ (longer recovery) |
| Sympathetic activation | ↓ (faster propagation) | ↓ (shorter recovery) |

### 2.3 Didactic Reentry Demonstration Mechanism

To make the concept of reentry observable in a classroom exercise, *iMyocyte* implements a structured demonstration that operationalizes Mines’ classical requirements for sustained reentrant excitation: a closed conduction pathway (ring), sufficient path length relative to refractoriness, and a substrate for unidirectional block [10]. In *iMyocyte*, reentry demonstrations use a closed ring topology in which each participant (cell) is connected to two neighbours (left/right), forming a circular conduction pathway. A designated leader assembles the ring by adding participants. The minimum ring size is 6 cells, ensuring that the circuit time (time for a wave to complete one loop) exceeds the refractory period of the unhealthy cell, thereby allowing the second wavefront to conduct through the recovered unhealthy cell and initiate reentry.

#### Initiation logic

After ring formation, the ring leader is to deliver a timed stimulation pulse that launches a bidirectional wavefront into the ring. Reentry is triggered deterministically when specific conditions are met at the unhealthy cell: if a wavefront reaches the unhealthy cell while it is refractory, the cell stores that wave’s unique identifier (UUID). If the same UUID returns after the refractory period has elapsed (i.e., the tissue has recovered), the unhealthy cell can re-excite and sustain circulating activity, producing a reentrant loop. This mechanism explicitly maps to the didactic point that reentry requires the returning wave to arrive after recovery, and that loop timing depends jointly on conduction velocity, refractory duration, and pathway length [3].

#### Interactive exploration of reentry conditions

To support active learning, the platform allows the ring leader to vary key factors that determine whether a circulating wave self-terminates, persists, or enters reentry (Figure 3). Cell health status is binary (healthy vs. a designated unhealthy cell), exercise level is set by the leader’s pacing of stimulation pulses (session-level), and sympathetic activation is a global on/off toggle applied uniformly across all cells. The leader’s UI also displays the reconstructed interaction graph (e.g., a ring topology) and displays each cell’s instantaneous CA state in real time, allowing wave propagation, refractoriness, and reentry dynamics to be monitored live. As such, users can visualize how:

- introducing an unhealthy cell creates a substrate for unidirectional block and re-excitation;
- increasing exercise level decreases propagation delay (*τ* ) and shortens the refractory period (RP), altering whether waves self-terminate or persist;
- enabling sympathetic activation produces the same qualitative shift (lower *τ*, shorter RP) across the ring.

**Figure 3:**
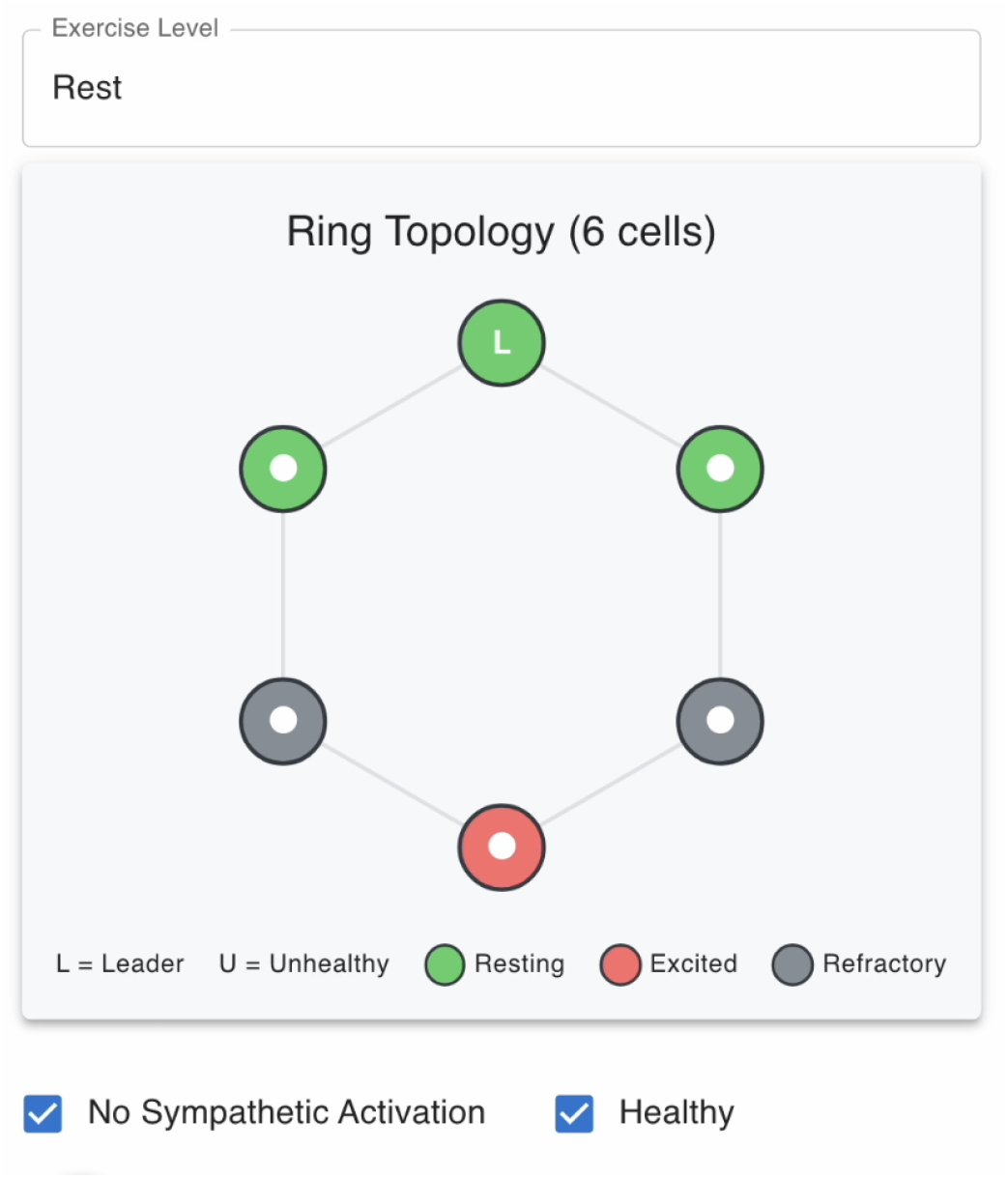
Leader interface snapshot during wave propagation in a ring topology. The UI displays the reconstructed ring (here, n = 6) and each cell’s instantaneous CA state in real time (legend shown in interface). In this frame, an excited wavefront (red) with a trailing refractory region (grey) is visible. For clarity, only the ring visualization and key controls are shown; sessionlevel controls allow the leader to set exercise level, toggle global sympathetic activation, and assign cell health status.

A screen-recorded demonstration of the leader UI during an active session is provided as Supplementary Video S1, showing stimulation, wave propagation, firing events, and induction of reentry after ring formation. Leader-triggered stimuli are used to illustrate reentry initiation (not continuous pacing).

#### Reentry termination

Once reentry is established, the demonstration can be terminated by the ring leader via a browser refresh action that clears the ring state and removes members from the topology. The system also automatically handles participant disconnection via Ably’s presence detection, dissolving the ring if the leader leaves or rebuilding the topology if other participants disconnect.

#### Intended learning objectives

After completing the *iMyocyte* reentry demonstration, learners should be able to: (i) describe how a closed pathway, unidirectional block, and recovery timing enable reentry; (ii) visualize how changing propagation delay (*τ* ) and refractory period (*T* ^ref^ ) shifts whether waves self-terminate or sustain; (iii) explain how heterogeneity (e.g., an unhealthy cell) alters loop timing and reentry susceptibility; and (iv) relate exercise/sympathetic modulation to qualitative changes in propagation and recovery.

### 2.4 Platform Verification

To validate correctness and cross-device synchronization, we implemented an automated integration test suite that exercises end-to-end behaviour across multiple clients under nominal network conditions. Tests covered:

- **State transitions:** Resting → Excited → Refractory → Resting; firing permitted only from Resting.
- **Timing parameters:** neighbour activations followed the implemented propagation-delay rule across representative parameter values (i.e., decreasing *τ*_*i*_ produced the expected qualitative increase in apparent wave speed), and cells enforced refractoriness such that restimulation attempts during 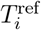 produced no excitation, whereas stimulation after 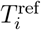 did.
- **Messaging:** stimulation events, parameter updates, and join/leave events were correctly applied to maintain session consistency.
- **Ring-topology checks:** verified correct assignment of two neighbours and event routing, including join/leave handling and topology updates.

### 2.5 Classroom Deployment and Evaluation Protocol

*iMyocyte* was deployed using a multi-phase evaluation strategy spanning (i) large-class piloting for feasibility and usability, and (ii) small-group testing for controlled observation of the reentry demonstration and targeted user feedback.

#### Phase 1: course pilot to assess baseline usability

A pre-release version of *iMyocyte* was piloted in an undergraduate classroom setting (McGill Physiology (PHGY) 210: Mammalian Physiology 2) to assess baseline usability and perceived pedagogical value in a real classroom setting. Feedback was collected through informal observation and post-session surveys, which identified key improvement targets (interface intuitiveness, waveform clarity, and simulation responsiveness).

#### Iterative refinement and re-deployment

Based on Phase 1 feedback, the platform underwent iterative refinement to improve interactivity, the visual representation of wave dynamics, and cross-device performance. The updated version was subsequently reintroduced in PHGY 210 as well as in a small-group demonstration designed for more structured input.

#### Phase 2: small-group reentry demonstration

A second round of testing was conducted with seven second-year undergraduate students (n = 7) from multiple life-science programs at McGill University. Participants used their own mobile devices and were assigned roles as individual cells within a closed ring topology; one participant was designated as the “unhealthy” cell to provide the pathological substrate required for reentry. The session was monitored using *iMyocyte*’s network interface, which visualizes the reconstructed ring topology and real-time cell-state updates.

#### Outcome measures

Evaluation emphasized practical, classroom-relevant endpoints: perceived ease of use, conceptual clarity, and perceived educational impact collected via post-session surveys, with qualitative comments used to guide further instructional and interface refinements.

## 3 Results

### 3.1 Platform Verification Supported Reliable Demonstrations

An automated integration test suite validated end-to-end behaviour across clients under nominal network conditions, including CA state transitions, timing parameter enforcement, event synchronization, and ring-topology routing. All verification tests passed, reducing the likelihood that the behaviours reported below reflect software or synchronization errors.

### 3.2 Initial PHGY 210 Deployment Indicated Improved Conceptual Understanding and Identified Usability Targets

In the first classroom pilot conducted in PHGY 210, post-session survey responses indicated that 57% of students felt *iMyocyte* “successfully” increased their understanding of the theoretical component, while 43% reported it “somewhat” increased understanding. Students also reported strong alignment with in-class content (61% selected the highest rating; 26% the next highest) and generally positive usability (48% selected the highest ease-of-use rating) (Figure 4). *iMyocyte* was consistently described as engaging, and qualitative feedback identified concrete goals for subsequent development, including (i) scaling to support more participants and longer-lived wave visualizations, and (ii) extending the platform to model other physiological/excitable systems beyond cardiac tissue.

**Figure 4:**
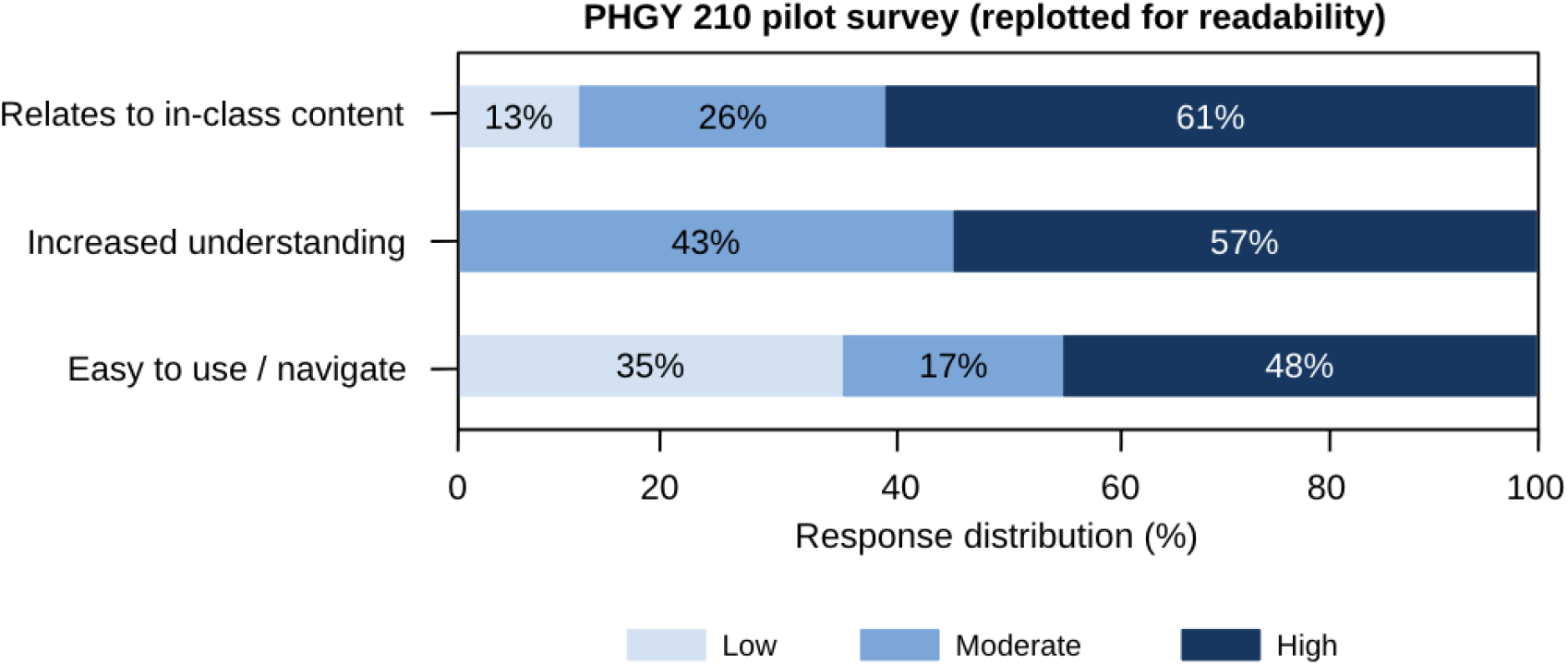
PHGY 210 pilot survey (replotted for readability). Responses were recorded on a 4-point ordinal scale (Lowest = 1, Low = 2, Moderate = 3, High = 4). No responses were recorded in category 1 (Lowest), which is omitted for readability. Percentages indicate the proportion of respondents in each category (n = 200).

### 3.3 Small Group Testing Reproduced Canonical Wave Termination versus Reentry in a Controlled Ring Topology

Following iterative refinement, a structured small-group session (n = 7) was conducted to enable controlled observation of wave dynamics in a closed loop. Screen recordings of *iMyocyte*’s network interface captured two distinct qualitative regimes consistent with excitable-media behaviour: in a healthy ring, an initial stimulus (S1) produced bidirectional propagation that self-terminated upon wavefront collision; in a ring containing an unhealthy cell, a second stimulus (S2) produced a transient conduction block, unidirectional propagation, and reentry after recovery, yielding sustained circulating activity (Figure 5).

**Figure 5:**
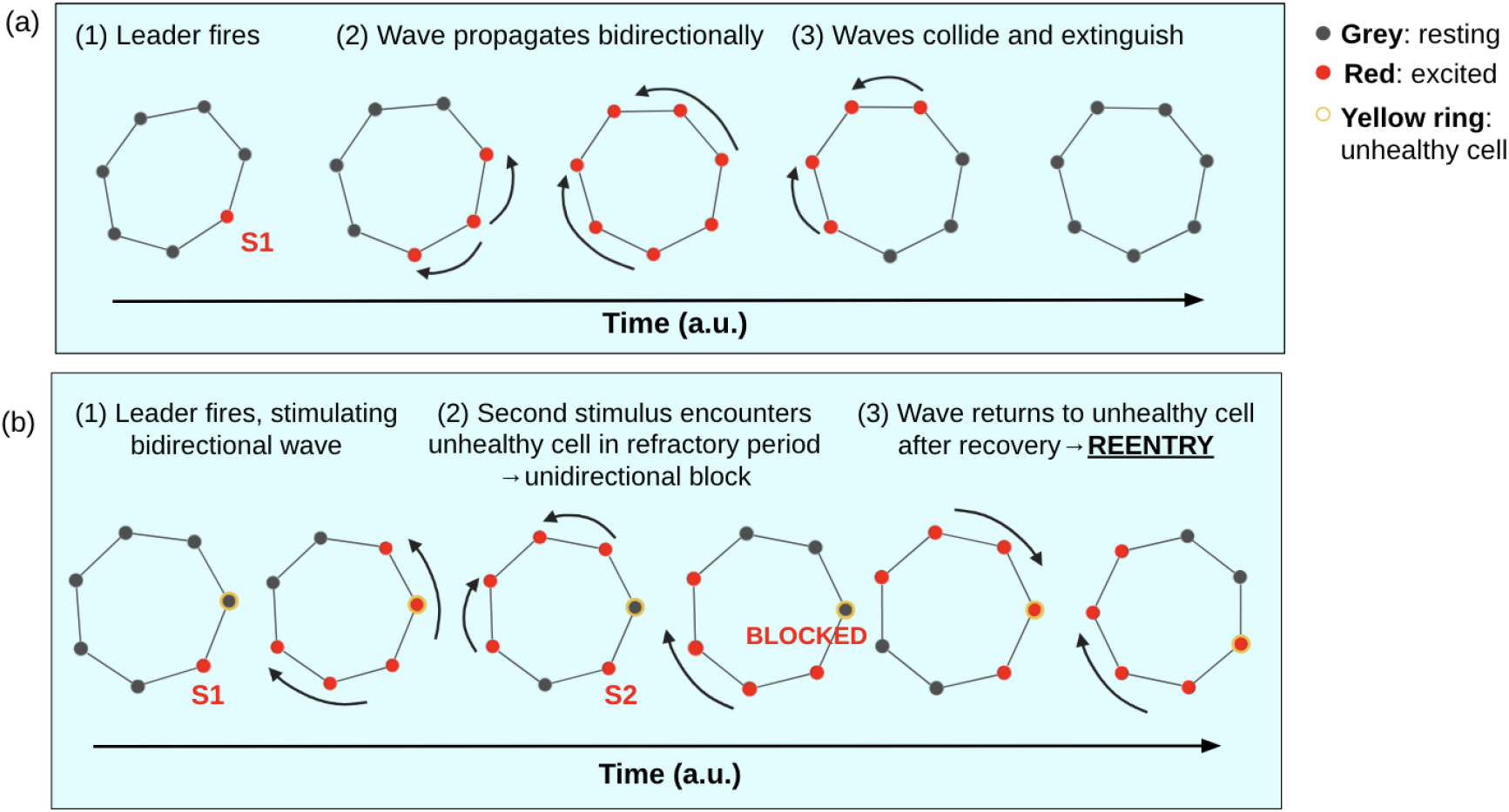
*iMyocyte* network-interface visualization of wave propagation from the n=7 pilot session. (A) Healthy ring: following an initial stimulus (S1), the excitation wave propagates bidirectionally and self-terminates when wave fronts collide. (B) Ring containing an unhealthy cell: following a second stimulus (S2), the wave encounters transient conduction block at the unhealthy cell, propagates uni-directionally, and re-enters after recovery, producing sustained circulating activity consistent with reentry.

In post-simulation surveys, all participants reported an improved understanding of the physiological conditions required for reentry, and specifically emphasized that stepwise manipulation of cell health states and external inputs clarified how action potential duration/refractoriness and conduction velocity interact to create arrhythmogenic timing conditions. For example, one participant noted that seeing the wave become blocked on one side “helped me finally understand how reentry happens not just in theory, but in motion.”

### 3.4 Large Classroom Deployment Revealed Scaling Failure Modes in Topology Formation and Signal Transmission

In a subsequent PHGY 210 demonstration involving almost 200 participants, multiple issues impacted the intended ring-based experiment. Although distinct rings were initially formed, these rings later merged into long chains, likely due to neighbour-selection errors that produced overlapping structures. Participant departures and the emergence of cells with more than two neighbours further destabilized the topology, enabling propagation between rings and disrupting expected patterns. One group exhibited reentry-like behaviour, but it was attributed more plausibly to signal transmission issues than to the intended deterministic trigger.

Despite these constraints, the deployment indicated that the updated client application supported real-time participation at scale in terms of interface operation and data collection. The session logs reported 189 unique cell IDs corresponding to 63 rings, with 697 total firing events. These logs confirm sustained classroom-scale connectivity and event exchange during the session, even when the intended ring topology was not preserved.

## 4 Discussion

*iMyocyte* demonstrates that a web-based, multi-user simulation can make core excitable-media concepts in cardiac electrophysiology tangible in real time. In small-group sessions, participants generated and observed qualitative regimes (e.g., wave propagation, conduction block, and reentry) while manipulating timing parameters (e.g., propagation delay, refractory duration) and introducing controlled heterogeneity within a ring topology, translating an abstract timing requirement into an observable phenomenon. More broadly, the platform’s decentralized design provides instructors with a flexible teaching format: the same interface can support intuitive demonstrations in small groups and, in principle, classroom-scale participatory experiments that emphasize emergent behaviour from local rules rather than memorization of definitions.

The primary limitations observed to date fall into two categories: (i) deployment and scaling constraints and (ii) modeling constraints. Although all integration tests passed, large-class deployments revealed scaling-related failure modes. In these settings, topology formation and maintenance became fragile: rings could merge into chains, participants could transiently acquire more than two neighbours, and network-level instability could disrupt intended propagation patterns. These failure modes reflect a real-world classroom setting (heterogeneous devices and variable connectivity) and motivate additional safeguards if ring-based demonstrations are to remain reproducible at ∼200-person scale.

Conceptually, *iMyocyte* is intentionally not a biophysically detailed tissue simulator. Its discrete, rule-based CA framework is designed to convey qualitative timing principles and therefore does not capture continuous voltage gradients, electrotonic spread, or finer-grained phenomena such as graded/partial conduction block, concepts relevant to reentry [20]. Accordingly, the platform should be interpreted as a didactic demonstrator rather than a quantitatively predictive electrophysiology model. Finally, while early surveys suggest high engagement and perceived clarity, the current evaluation is not a controlled learning study and does not yet quantify learning gains relative to standard instruction.

Several upgrades would directly address the observed scaling failure modes and strengthen reproducibility: (i) Topology hardening by locking ring topology once a demonstration starts and enforcing a strict two-neighbour cap in ring mode. (ii) Stronger synchronization, including explicit acknowledgements for critical events, leader-driven resynchronization checkpoints, and improved handling of late or duplicate messages) to reduce jitter and divergence across devices. (iii) Formal pedagogical evaluation using pre/post concept assessments, comparison against standard instruction, and retention testing to convert perceived clarity into measurable learning outcomes.

Overall, *iMyocyte* provides an accessible, interactive, and collaborative way to teach the timing logic that underlies conduction block and reentry. In small-group settings, the platform reliably supported hands-on exploration of how conduction velocity and recovery interact to permit or prevent sustained circulating activity, aligning with its goal as a classroom demonstrator. At larger scales, topology instability and variable message timing point to two engineering priorities: enforcing ring invariants and improving synchronization, which are necessary for robust demonstrations in very large cohorts. With these refinements and formal learning assessment, *iMyocyte* could serve as a generalizable framework for participatory teaching of excitable-media dynamics, turning classrooms into distributed, interactive modeling environments.

## Acknowledgements

I would like to thank McGill undergraduate students Emma Neamtu, Athena Chen, Emile Fontaine, Hannah Zheng, Siddharth Gollapudi, Elena Pivovarcic, Zaina Abodan, and Salim Tarbouche for volunteering as test participants and providing invaluable feedback during the small-group pilot study.

